# Half of the world’s tree biodiversity is unprotected and is increasingly threatened by human activities

**DOI:** 10.1101/2020.04.21.052464

**Authors:** Wen-Yong Guo, Josep M. Serra-Diaz, Franziska Schrodt, Wolf L. Eiserhardt, Brian S. Maitner, Cory Merow, Cyrille Violle, Madhur Anand, Michaël Belluau, Hans Henrik Bruun, Chaeho Byun, Jane A. Catford, Bruno E. L. Cerabolini, Eduardo Chacón-Madrigal, Daniela Ciccarelli, Johannes H. C. Cornelissen, Anh Tuan Dang-Le, Angel de Frutos, Arildo S. Dias, Aelton B. Giroldo, Kun Guo, Alvaro G. Gutiérrez, Wesley Hattingh, Tianhua He, Peter Hietz, Nate Hough-Snee, Steven Jansen, Jens Kattge, Tamir Klein, Benjamin Komac, Nathan Kraft, Koen Kramer, Sandra Lavorel, Christopher H. Lusk, Adam R. Martin, Maurizio Mencuccini, Sean T. Michaletz, Vanessa Minden, Akira S. Mori, Ülo Niinemets, Yusuke Onoda, Renske E. Onstein, Josep Peñuelas, Valério D. Pillar, Jan Pisek, Bjorn J.M. Robroek, Brandon Schamp, Martjin Slot, Ênio Sosinski, Nadejda A. Soudzilovskaia, Nelson Thiffault, Peter van Bodegom, Fons van der Plas, Ian J. Wright, Wu-Bing Xu, Jingming Zheng, Brian J. Enquist, Jens-Christian Svenning

**Author notes:** (WYG); (WBX); (JCS); (WE), (BSM); (BJE), (AdF); (REO),;, (NS); (PMvB).

## Abstract

Although trees are key to ecosystem functioning, many forests and tree species across the globe face strong threats. Preserving areas of high biodiversity is a core priority for conservation; however, different dimensions of biodiversity and varied conservation targets make it difficult to respond effectively to this challenge. Here, we (*i*) identify priority areas for global tree conservation using comprehensive coverage of tree diversity based on taxonomy, phylogeny, and functional traits; and (*ii*) compare these findings to existing protected areas and global biodiversity conservation frameworks. We find that *ca*. 51% of the top-priority areas for tree biodiversity are located in current protected areas. The remaining half top-priority areas are subject to moderate to high human pressures, indicating conservation actions are needed to mitigate these human impacts. Our findings emphasize the effectiveness of using tree conservation priority areas for future global conservation planning.

## Introduction

Trees play a vital role in the Earth system as the principal agents of energy and matter in terrestrial ecosystems. The diversity and functioning of trees structure biological diversity and provide multiple ecosystem services, with large trees playing an especially vital role in governing rates of ecosystem services such as catchment and watercourse protection, carbon sequestration, and climate regulation (*1–3*). Trees provide habitat for many other taxa (*4–8*). The magnitude of many of these functions and services is generally expected to increase with as tree diversity and abundance increase. The functional diversity of tree assemblages enhances ecosystem productivity and stability (*9–11*). However, continued deforestation (*12–17*) results in biodiversity loss of both tree species themselves and tree-dependent organisms (*8, 18, 19*), decreases in ecosystem productivity (*20*), and also jeopardizes other ecosystem functions such as climate regulation (*21, 22*). Consequently, it is urgent to improve the conservation of high-diversity areas for trees to protect not just tree diversity, but also associated biota, the ecosystems in which they occur, and the processes and services that these ecosystems provide (*23*).

Protected areas (PAs) are the primary conservation strategy for preventing biodiversity loss and preserving nature (*24–26*). In recent decades, the expansion of PAs has been considerable, with ∼15% of the Earth’s land now located within PAs (the World Database on Protected Areas, WDPA; https://livereport.protectedplanet.net/chapter-2, accessed on April 15, 2020). This coverage is close to the Convention on Biological Diversity (CBD) 2020 target of protecting at least 17% of Earth’s land area. Despite the rapid progress of PA expansion worldwide, research has found that current PAs do not overlap with certain regions that harbor otherwise high taxonomic diversity (e.g., mammals (*27, 28*). Additionally, approximately one third of global PAs are experiencing intense human pressure (*29*), and this phenomenon is likely to continue globally as human activity increases near existing protected areas (*26, 30*).

To halt or slow continuing biodiversity declines, the post-2020 protected area targets are being intensely debated (*31–33*). One initiative that has gained significant momentum is Wilson’s “Half Earth” proposal (*34*), which affirms that half of the Earth’s surface needs to be protected in order to safeguard important areas of biodiversity and ecosystem processes and services (*35–37*). This suggestion is also in accordance with the CBD’s proposed 2050 goal related to its Post-2020 Biodiversity Framework (https://www.cbd.int/conferences/post2020/post2020-prep-01/documents, accessed on March 20, 2020). While global conservation targets and locations are negotiated by intergovernmental organizations (*38*), the proposed protected locations should be representative areas with high levels of biodiversity, endemics and ecosystem services. Indeed, the emphasis on increased biodiversity and ecosystem services can be used to test the effectiveness of existing PAs. Given the vital role of tree species to Earth systems, tree diversity is a practical indicator to the effectiveness of the existing PAs and predictor to allocate protected targets for CBD 2020 and beyond (e.g. CBD 2050). However, neither the effectiveness or predictors to allocate future targets were previously investigated, nor the extent of the tree diversity protection.

Here, we assembled a novel global database to identify priority areas for conservation of tree diversity worldwide. Following (*39*), our analyses use the Zonation method (*40–42*) and integrates multiple measures of diversity including taxonomic distribution data, phylogenetic information, and functional trait data. We then assessed whether or not priority areas for tree conservation overlapped or diverged spatially by estimating the proportions of overlap among and between the top 17% (i.e., CBD 2020 target) and 50% (i.e., CBD 2050 target) priority areas across three dimensions of diversity: taxonomic, phylogenetic, and functional diversity (*27, 43*). Additionally, we selected three existing global biodiversity conservation priority frameworks (the Global 200 Ecoregions (G200), the Biodiversity Hotspots (BH), and Last of the Wild (LW), Brooks *et al*., 2006) proposed by certain non-governmental organizations (NGOs), and compared the spatial overlap between the top priority areas, existing PAs obtained from WDPA, and the selected conservation priority frameworks. We made this comparison to identify gaps and overlaps between spatial prioritization outcomes, current conservation efforts, and existing global conservation frameworks. Moreover, we compared the intensity of human pressure using a human modification index (*44*) (a cumulative measure of human alteration of terrestrial lands) inside and outside current PAs for the areas of the prioritization outcomes. Based on multiple facets of tree species diversity extracted from the most extensive global tree species distributional database, a dated phylogeny, and eight ecologically important functional traits, we assess the quality of current PAs and identify and evaluate potential future locations for PA expansion.

## Results

### Global priority areas for tree conservation across the three dimensions of diversity

Priority areas for tree conservation diverged spatially among taxonomic, phylogenetic, and functional diversity metrics. This was more evident in the case of the CBD 2020 target areas (top 17% priority) than of the CBD 2050 target areas (top 50% priority) (Figs. 1 & S1). The areas that were prioritized for all three dimensions occurred primarily in the tropical rainforest regions of the Americas, Africa, Indo-Malaya, and Australasia, as well as in subtropical Asia. Generally, areas prioritized by just two diversity metrics were adjacent to these areas, such as the inland regions of Brazil and Australia (Fig. 1a). High priority sites for tree conservation for any given single diversity dimension were scattered globally (Fig. 1a). Although nearly half of the 2020 target areas (8.2% in the top 17%) were shared by the priority areas for all three tree diversity dimensions, about 7.4% of the 2020 target areas based on taxonomic diversity differed from those based on phylogenetic and functional diversity (Fig. 1b). In contrast, the top 17% phylogeny- and trait-based priority areas largely overlapped, reflecting high correlation of phylogenetic and functional diversity in this study (Fig. 1a & 1b, Table S1).

**Fig. 1.**
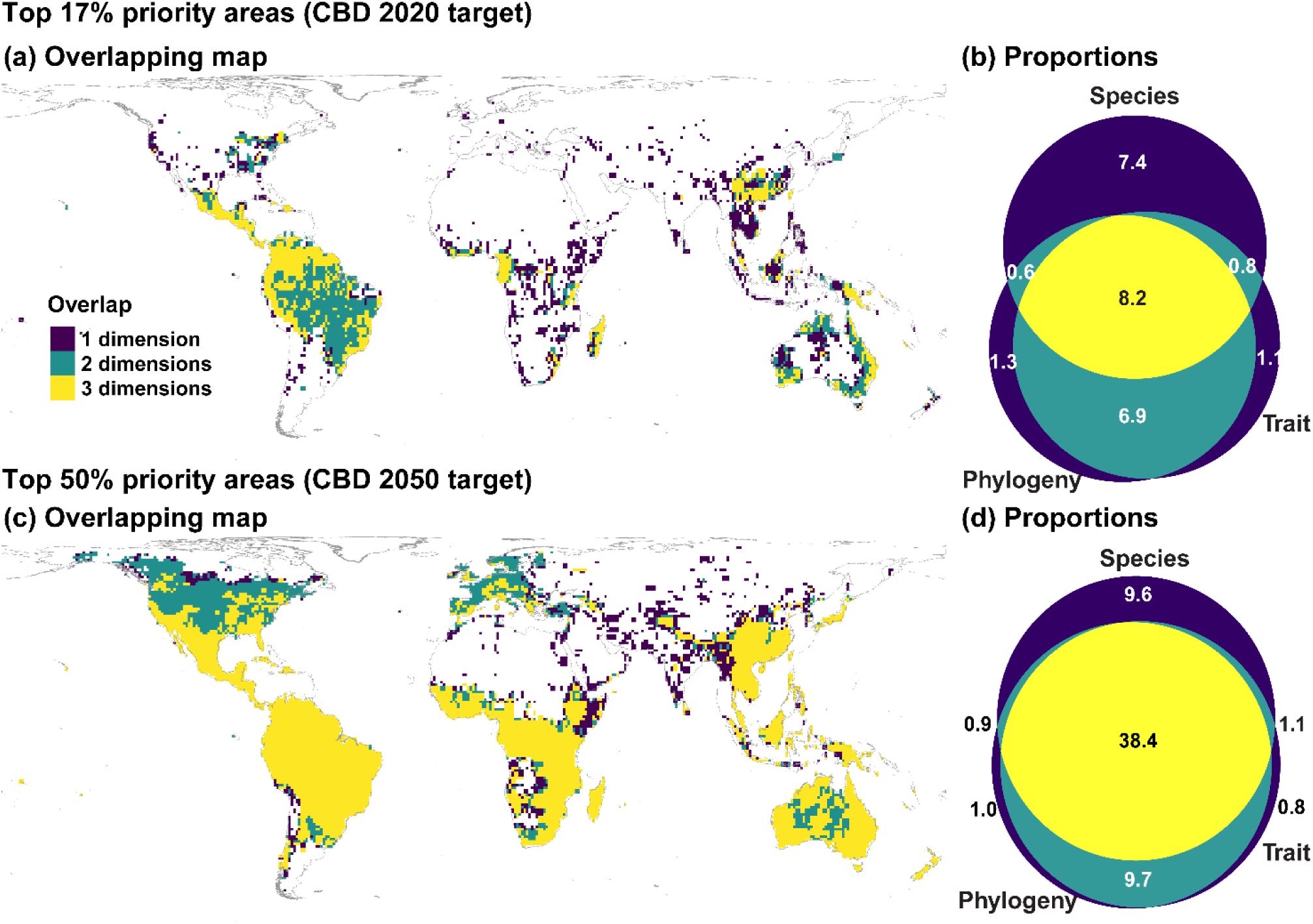
The top 17% (CBD 2020 target, upper panel) and 50% (CBD 2050 target, lower panel) priority cells (a, c) using species, phylogenetic and functional diversity facet according to zonation prioritization; (b, d) Venn diagrams showing the proportion of the land surface with overlap between species, phylogenetic and functional dimensions of diversity for the top 17% and top 50% priority areas. Colors indicate the overlaps between combinations of two of the two facets of the three facets, between all three facets, or no overlap.

When extending the priority areas to the 2050 target 2050, most areas (38.4% in the top 50%) were shared by all three dimensions (Fig. 1d), and located primarily in the tropical and subtropical regions, as well as in some temperate regions (Fig. 1c). Relatively large temperate areas were only prioritized when evaluating phylogenetic- and functional trait diversity (Fig. 1c).

### Congruence between top tree conservation priority areas, existing PAs, and each conservation priority framework

About half of the 2020 or 2050 target areas for tree priority conservation (jointly optimized in taxonomic, phylogenetic, and functional diversity) are currently protected in existing conservation sites (51.2% and 44.6% respectively, the proportion of “WDPA only” and “Shared” in Fig. 2a & 2b). For the three conservation priority frameworks (Fig. S2), the G200 always had higher overlap with top priority areas for tree diversity (45.4%), the BH the second (34.8%), and LW the least (7.3%) (Fig. 2a). About 91% of the 2020 target areas would be protected if PA expansion is based on the G200 framework, while 77% would be protected for the 2050 target areas. On the other hand, only about 50% of the top priority areas (56% for 2020 and 52% for 2050 targets) would be protected if LW is used to govern future conservation planning. Providing an intermediate case, about 73% of the 2020 target and 66% of the 2050 target areas would be protected if the BH framework were to be used as the guideline.

**Fig. 2.**
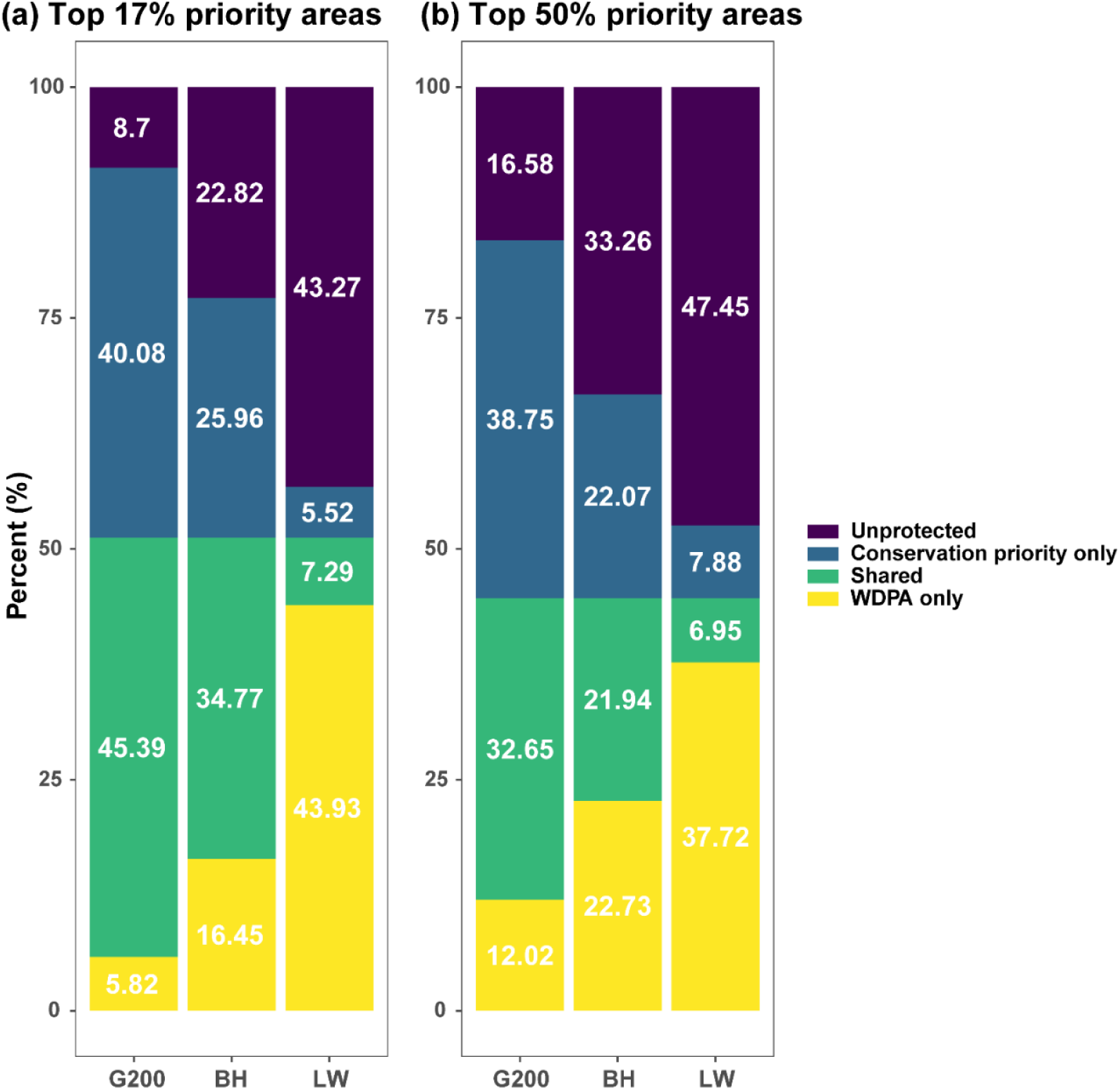
Percentages of the top 17% (CBD 2020 target) and top 50% (CBD 2050 target) priority areas for tree diversity (jointly considering taxonomic, phylogenetic, and functional diversity) covered by the existing protected areas (WDPA) or by each global biodiversity conservation priority framework (G200, BH, and LW). Colors indicate the overlaps between combinations, Unprotected: areas not overlapping with either WDPA or the conservation priority framework; Conservation priority only: areas overlapping with conservation priority framework only; Shared: areas overlapping with both WDPA and conservation priority framework; WDPA only: areas overlapping with WPDA only. G200: Global 200 Ecoregions; BH: Biodiversity Hotspots; LW: Last of the Wild. The tree conservation priority areas were obtained by using the three diversity dimensions simultaneously in Zonation.

### Human pressure of tree priority areas, inside and outside existing PAs

Most of the top 17% tree conservation priority areas suffered moderate human pressure with human modification values between 0.1 and 0.4, such as in southern and eastern Asia, South America outside the Amazon Basin, and Madagascar (Fig. 3). On the contrary, many of the top 50% priority areas were subject to high human pressure, with human modification values greater than 0.4 (blue and red regions in Fig. 3). These areas included many European countries, India, eastern China, Indonesia, Nigeria, Ethiopia, central North America, and eastern Argentina.

**Fig. 3.**
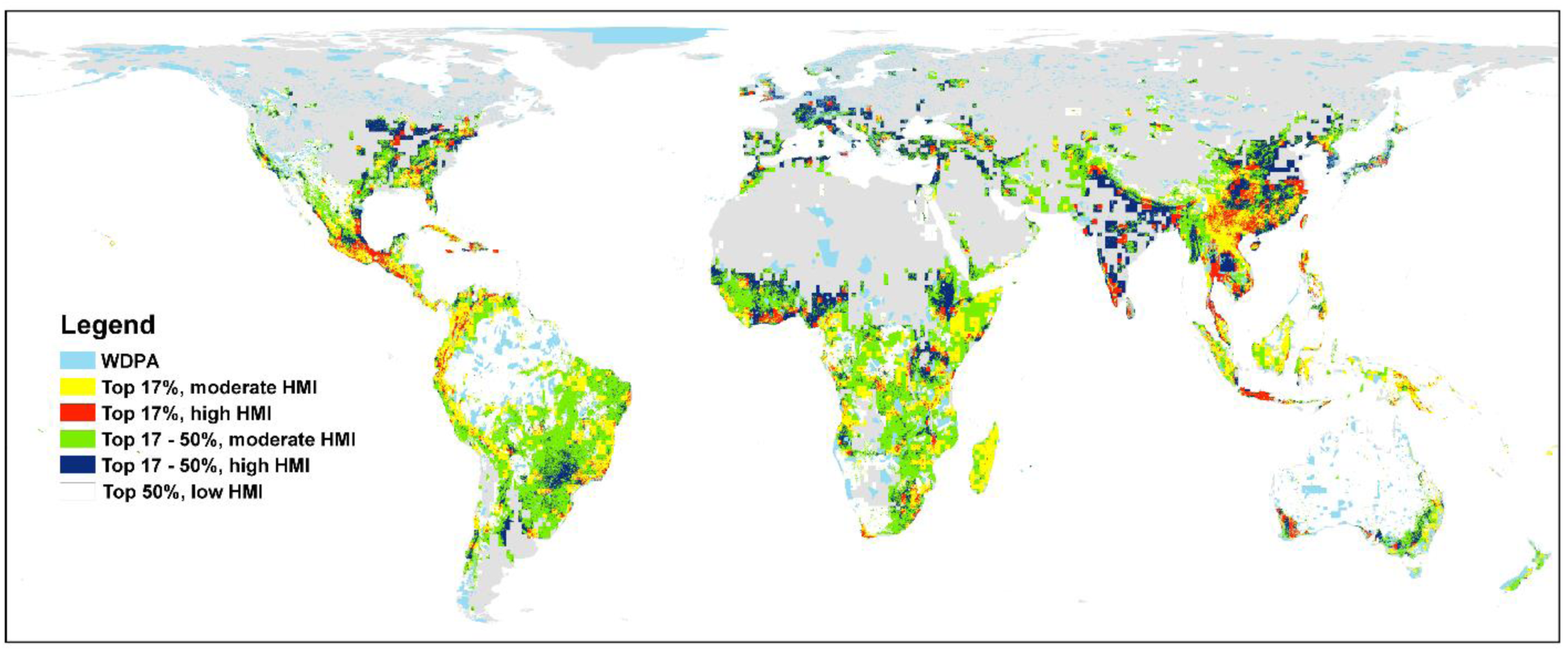
Congruence between current protected areas (PAs, following the World Database on Protected Areas (WDPA) database), top priority areas for tree diversity conservation (jointly considering taxonomic, phylogenetic, and functional diversity), and moderate to highly human-modified land cover. The categories of the human modification index (HMI) represent low (≤0.1), moderately (0.1 - 0.4) and highly to very highly (>0.4) modified land cover, respectively (Kennedy et al., 2019). The WDPA layer is the polygons of the current PAs. The human modification index layer is shown in a resolution of 1-km^2^.

The global average human modification index was lower in the areas inside PAs than in those outside PAs (Fig. S3, mean values of 0.07 vs. 0.19, one-way ANOVA, *p* < 0.0001). The mean human modification values inside the top 17% and top 17-50% priority areas were similar (Fig. 4, 0.10 cf. 0.08), and the same to outside PAs (Fig. 4, 0.25 cf. 0.24), while for both priority areas the mean human modification value inside and outside PAs significantly differed (*p* < 0.0001, Fig. 4). Mean human modification values were higher inside PAs than the corresponding global mean human modification values (*p* < 0.0001).

**Fig. 4.**
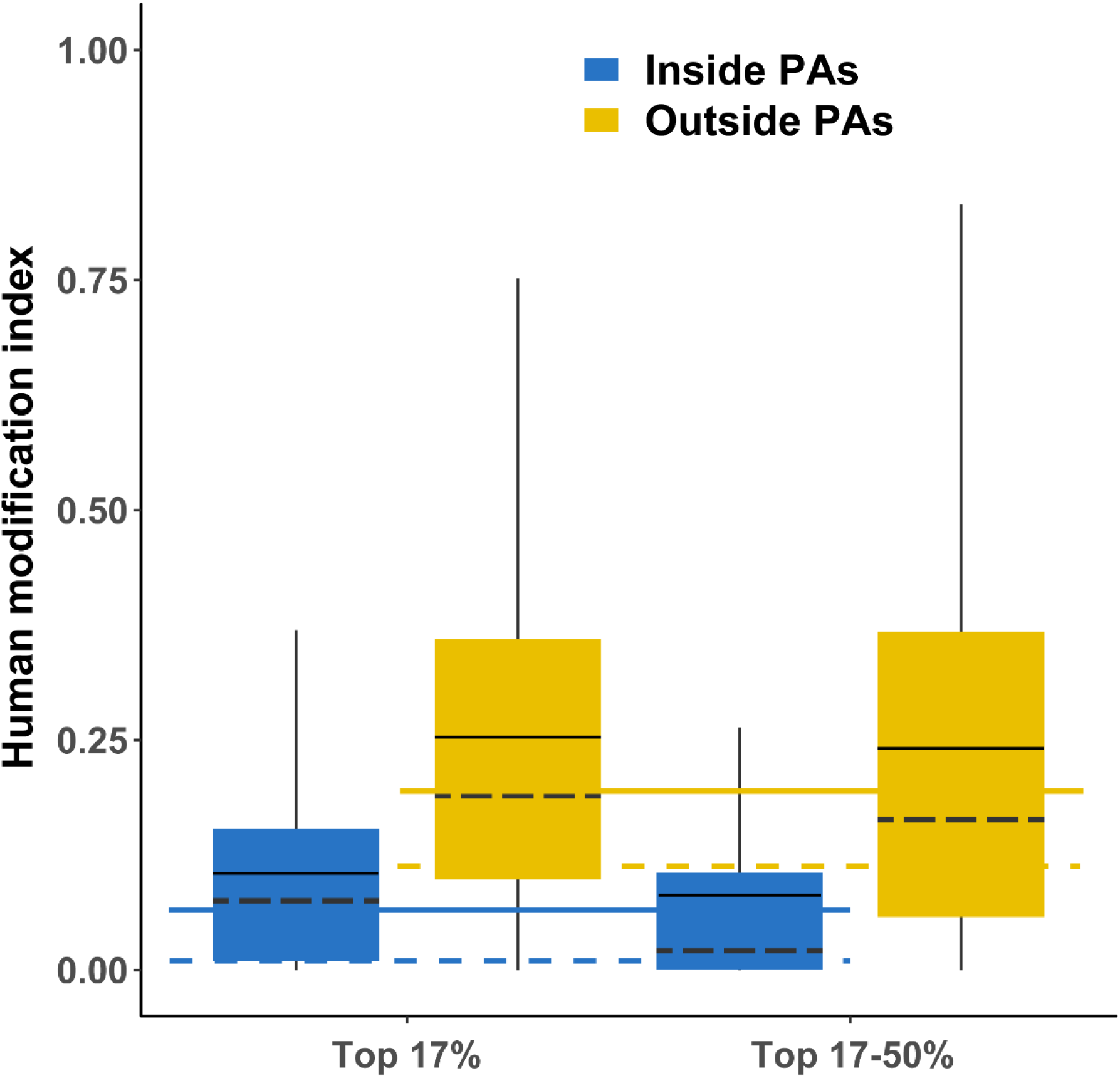
Boxplot of the human modification index across tree diversity conservation categories (top 17%, and top 17 - 50% of the priority scores from the Zonation prioritization jointly considering taxonomic, phylogenetic, and functional diversity). Each category is divided into two groups: areas inside (blue) and outside (orange) protected areas (PAs, following the World Database on Protected Areas (WDPA) database. Medians and means each are indicated by solid lines and dashed lines in the boxplots. The horizontal solid lines and dashed lines indicate the global median and mean human modification values for the areas inside and outside PAs, respectively. All values were obtained from 1-km^2^ resolution input layers (see Methods for more information).

## Discussion

Given the importance of tree diversity as a component of Earth’s biodiversity, for the functioning of forest ecosystems, and for the multiple ecosystem services, conservation and restoration of high conservation priority for tree diversity is critical. Using multiple dimensions of diversity, we identified areas of high conservation importance across the world. Currently, only about half of the most important tree biodiversity conservation priority areas are located inside PAs, which are defined by less intense human activities (e.g., the Amazon Basin and middle-west North America). However, the other half of high-priority areas for tree diversity conservation are distributed in regions with moderate to high human pressures, where certain kinds of threats, e.g., fire, habitat conversion, or overgrazing, are likely to negatively impact ecosystems (*16, 17, 45, 46*). Thus passive restoration, such as natural regeneration on degraded lands, is likely to be a more effective and affordable approach to recover and conserve the ecosystem structure and function (*47, 48*), alongside using legal boundaries and designations to protect land from human pressures (*49*). Schemes for conservation of biodiversity outside of PAs (e.g. protection under Payments of Ecosystem Services, PES) could also be sponsored by governments, NGOs and companies with social responsibility as a tool to preserve biodiversity and reduce the human pressure on these areas (*50*).

The high overlap between the conservation priority frameworks and the top-priority areas for tree diversity, particularly the Global 200 Ecoregions and the Biodiversity Hotspots frameworks, pinpointed the representativeness of tree diversity to a broader set of organisms, as these frameworks selected ecoregions of most crucial (either high irreplaceability or vulnerability) to the global biodiversity (*51, 52*). On the contrary, the Last of the Wild framework had the worst match with the tree conservation priority areas, primarily because this conservation scheme primarily prioritizes areas with climates less suitable for tree growth, e.g., arid deserts and cold high altitudes or latitudes in the Northern Hemisphere, where tree species diversity is inherently low (*53*). However, these areas have also incurred less overall human pressure (Figs. 3 & 4), making conservation implementation less costly (*25, 29, 31, 54*).

The tree conservation priority areas based on three dimensions of biodiversity are spatially incongruent (Fig. S1), particularly between taxonomic diversity on one hand and phylogenetic and functional diversity on the other, indicating the importance of considering multiple biodiversity dimensions in conservation planning (*27, 43*). Previous conservation priority studies have frequently used just one dimension, typically taxonomic diversity (*55–57*), as a surrogate for other aspects of biodiversity and ecosystem function (*28, 58, 59*), and sometimes argued that incorporating phylogenetic and functional data would not be necessary.However, each diversity facet represents important ecological and evolutionary elements of biodiversity, and many studies report spatial mismatches among these dimensions (*28, 60–64*), as we found for tree diversity globally. We found the non-taxonomic prioritization analysis could provide *ca*. 10% more high priority areas for the top 17% and 50% priority areas (Fig. 1). More importantly, the priority areas defined by phylogenetic and functional traits are more spatially continuous than those determined by species richness (similar as (*27*)). Given current conservation planning activities tend to focus on large and highly connected areas (*24, 34, 65*), the incorporation of phylogeny and functional trait dimensions could provide additional insights for conservation planning.

## Conclusions

With the end of the Decade on Biodiversity, a new UN Decade on Ecosystem Restoration starts from 2021. Knowing where to protect and restore biodiversity is imperative to support conservation activities. Considering the global diversity of trees, we show that about half of the critical sites for tree diversity are well protected with little human disturbance. Conversely, the remaining half of these priority areas is not protected and largely located in places with high anthropogenic activities, thus, future conservation planning should be more based on restoration actions. Meanwhile, attention should also be paid to the increasing pressure within the existing protected areas (*26, 29, 38*), as rising human pressure is a risk in many places (e.g., the Serengeti-Mara ecosystem (*66*)), in addition to rising pressure from human-driven climate change (*67*) or extreme disturbance events such as mega-fires (*68*). From the aspect of tree biodiversity, the existing conservation priority frameworks are helpful for planning and implementing future PAs, especially the Global 200 Ecoregion framework, which provides the best overlap with priority areas for global tree diversity conservation.

## Methods

### Tree species and their occurrence records

In this study, we used the world tree species list and species occurrence data compiled and cleaned by (*69*). Briefly, they extracted the records from the world tree species checklist (GlobalTreeSearch, GTS; (*70*)), and further standardized the taxonomic names via the Taxonomic Name Resolution Service (TNRS) online tool (*71*). The GTS employed the definition of the tree type growth habit agreed by IUCN’s Global Tree Specialist Group (GTSG), i.e. “a woody plant with usually a single stem growing to a height of at least two meters, or if multi-stemmed, then at least one vertical stem five centimeters in diameter at breast height” (*70*). A dataset of 58,100 tree species was obtained (*69*).

They further collected the occurrence data for the tree species (*69*). Five widely used and publicly accessible occurrence databases were used: the Global Biodiversity Information Facility (GBIF; http://www.gbif.org), the public domain Botanical Information and Ecological Network v.3 (BIEN; http://bien.nceas.ucsb.edu/bien/; (*72, 73*)), Latin American Seasonally Dry Tropical Forest Floristic Network (DRYFLOR; http://www.dryflor.info/; (*74*)), RAINBIO database (http://rainbio.cesab.org/; (*75*)), and the Atlas of Living Australia (ALA; http://www.ala.org.au/). Due to well-documented problems of biases and errors in global plant occurrences datasets (*76*), a workflow was further developed for occurrence data quality assessment and control, and successfully labelled all the occurrence data with a quality level (*69*). We initially had 9,032,654 non-quality assessed occurrence records. We adapted the quality category of the occurrence records in the study, and selected occurrence records with quality labels corresponding to AAA, AA, A, and C. These correspond to data not being an outlier in the environmental space, and not in urban areas or in botanical gardens. Our final list of species was lowered to 46,752 species with a total occurrence dataset of 7,066,785 records. Although we obtained a relatively comprehensively geographical coverage of the tree species occurrence data (*69*), some regional gaps still present, such as Russia. We thus downloaded the distribution range data of 44 species from the European Forest Genetic Resources Program (EUFORGEN, http://www.euforgen.org/) to fill the coverage gaps in Eastern Europe and Russia.

### Species range estimates and external validation

We constructed alpha hulls to estimate the range of each species with 20 or more occurrence records using the “*ashape*” function of the *Alphahull* package (*77*) implemented in R (ver. 3.5.1; (*78*)), which is based on the algorithm by (*79*). For species with less than 20 occurrences or with disjunct records, a 10-km buffer was given to each point record and then combined with the alpha-hull range. Previously, several alpha levels were recommended for the estimation of species range (e.g., (*80–83*)), four alpha levels (2, 4, 6, 10) were applied to each species here. The obtained estimated range maps were rasterized to 110 km equal-area grid cells, a resolution commonly used in global diversity studies (e.g., (*84–86*)), using the *letsR* package (*87*).

To validate the range maps using different alpha levels, we performed an external validation against the results of an independent modelling study published by (*53*). In their study, the authors used cross-scale models to predict the tree species richness by integrating 1,336 global forest plots and 282 regional checklists. They produced two spatial levels of richness maps: one in one ha plots and one in 209,903 km^2^ hexagons. Here we compared our data with their coarse-grained predicted richness maps only. By following a similar method (*53*), we firstly calculated the species richness in each 209,903 km^2^ hexagon for each of the alpha-level range maps. We then plotted the observed richness in these hexagons against the predicted richness in ref (*53*). Although the range extents from the four alpha levels were different, the species richness obtained from them were largely consistent, as indicated from the similar intercepts and slopes of the regression lines between the predicted species richness from ref (*53*) and observed species richness of each alpha-level range map, although the predicted value of species richness per hexagon were generally higher than those from alpha-hull estimations (Figs. S4-S5). In fact, two of the three datasets ref (*53*) used for external validation were the same as ours, i.e., BIEN and RAINBIO. We selected the alpha-hull range maps with an alpha parameter of 6 for subsequent analyses, as it is commonly used (*82, 83*).

### Phylogeny

We extracted phylogenetic information for the tree species with range maps from the largest seed plant phylogeny that is currently available (the ALLMB tree (*88*)). This phylogeny combines a backbone tree (*89*) reflecting deep relationships with sequence data from public repositories (GenBank) and previous knowledge of phylogenetic relationships and species names from the Open Tree of Life (Open Tree of Life synthetic tree release 9.1 and taxonomy version 3, https://tree.opentreeoflife.org/about/synthesis-release/v9.1). We matched this phylogeny to our tree dataset by first removing any species that are not in our data, and then manually adding some species that were missing from the phylogeny (due to different taxonomic concepts) following the same approach that ref (*88*) used to add missing species. The resulting phylogeny contained 46,752 species (Fig. S6).

To represent the phylogenetic position of each species in our dataset, we calculated phylogenetic eigenvectors (*90*) using the *PVR* package (*90, 91*). Because we were mostly interested in the deep structure of the phylogeny, and phylogenetic eigenvector calculation for large phylogenies is computationally prohibitive, we calculated eigenvectors at the genus level (4,031 genera). To accomplish this, we randomly chose one species per genus, removed all other species, and computed phylogenetic eigenvectors using the resulting phylogenetic tree. All species were then assigned the eigenvector values of their genus. Because some genera were not monophyletic in the tree (largely due to lack of resolution), we repeated this process 800 times to capture phylogenetic uncertainty, and averaged resulting eigenvector values across the 800 replicates. In the following analysis we used the first 15 eigenvectors, excluding those that captured very little phylogenetic variation (eigenvalues <1%, following (*27*)). These selected eigenvectors accounted for 40.6% of the total phylogenetic variation, representing the deep evolutionary history of our study species.

Phylogenetic eigenvectors represent phylogenetic placement as continuous variables, but our subsequent analysis required a binary presence-absence matrix. To accommodate this, we adapted from the framework of ref (*27, 43*). First, we evenly divided each eigenvector into 20 bins. Then we created a binary variable for each bin, scoring all species with values within the range of the bin 1, and all others 0. This resulted in 20 binary variables for each of the 15 eigenvectors, i.e. a matrix of 46,752 species × 300 binary variables. We multiplied this matrix with the grid cells × species matrix to generate a presence-absence matrix of phylogenetic groups in grid cells (*27*). This matrix showed which parts of the phylogeny (as represented by the binary variables derived from the eigenvectors) were present in each grid cell. We used this matrix in the following prioritization analysis to find priority regions for the conservation of tree phylogenetic diversity.

### Functional trait data

Twenty-one functional traits (Table S2) were compiled using the TRY (https://try-db.org/TryWeb/Home.php; (*92, 93*)), TOPIC (*94–100*) and BIEN (http://bien.nceas.ucsb.edu/bien/; (*72*)) databases. As many of the traits had missing values (more than 88% of the data per trait), we applied gap-filling with Bayesian Hierarchical Probabilistic Matrix Factorization (BHPMF (*101–103*)), which is a robust machine learning gap-filling technique with the consideration of phylogenetic trait signal and trait–trait correlations, and has been successfully applied in recent global trait-related studies (e.g., (*104, 105*)). To increase the imputation accuracy, we included phylogenetic information in the form of the phylogenetic eigenvectors (PEs) obtained from phylogenetic eigenvector analysis, as suggested by (*106*). As the inclusion of PEs greatly changed the gap-filling results, we ran several trials, and found the Root Mean Squared Error (RMSE) was the smallest when the first six PEs were included in the imputation (Fig. S7). We thus used the first six PEs in the final gap-filling process. We further used the minimum and maximum values per trait of the observed data as thresholds. If the gap-filled data were outside of the thresholds, the observed minimum or maximum were used to replace the gap-filled data. As for plant maximum height, we further used a height of 2 m to replace any imputed data lower than that, after the definition of tree species used in the GlobalTreeSearch database (*70*). We finally selected eight ecologically relevant and common-used traits (*104*) for further functional diversity analyses, that is, leaf nitrogen content, wood density, leaf phosphorus content, leaf dry matter content, plant max height, seed dry mass, specific leaf area, and leaf area (*107*). We used the *beanplot* package (*108*) to visually compare the observed original and imputed data, and found they generally had similar distribution patterns for each functional trait (Fig. S8).

As all the eight functional traits are continuous, to obtain trait group distribution maps we followed a similar procedure as in the case of phylogenetic eigenvectors. Briefly, we split each trait into 20 equal bins, and then converted it into a binary species × trait matrix for each trait (46,752 × 20). For each of the 20 bins, we multiplied it by the 110 × 110 km grid cells × species matrix to obtain a trait × grid cells matrix, in which each 110 × 110 km grid cell contained the number of species of trait values within the trait interval. Totally, we obtained 160 trait × grid cells matrices, a distribution map was generated for each of them, and then all the 160 distribution maps were used in the prioritization analysis to locate the priority regions for trait dimension.

### Protected area data

The protected area (PA) data were extracted from the December 2019 release of the World Database on Protected areas (WDPA) via the *wdpar* package (*109, 110*). The release includes 244,869 PAs globally. According to previous similar global studies (e.g., (*29*)), we extracted the protected areas from the WDPA database by selecting terrestrial areas belonging to IUCN protected area categories I to VI and having a status “designated”, “inscribed”, or “established”, and areas not designated as UNESCO Man and Biosphere Reserves. We also excluded the PAs represented as points. Totally 95,506 PAs were kept and then resampled at the 110 × 110 km grid cell level.

### Global biodiversity conservation priority frameworks

Many NGOs have proposed frameworks for global biodiversity conservation prioritization, e.g., the Biodiversity Hotspots (BH) by Conservation International (*51*), the Last of the Wild (LW) by the Wildlife Conservation Institute (*111*), and the Global 200 Ecoregions (G200) by the World Wide Fund For Nature (*52*). However, these frameworks vary in both location and coverage, largely due to the different emphasis of the nature conservation organizations. Although the two central aspects of systematic conservation planning, irreplaceability and vulnerability (*24*), are equally important, some of the conservation organizations concentrate only on irreplaceability, while others focus more on vulnerability (*25*). Under the framework of irreplaceability and vulnerability, Brooks et al. (*25*) summarized nine major frameworks, dividing them into three groups: prioritizing high vulnerability (regions of high threat, purely reactive, e.g., BH), low vulnerability (regions of low threat, purely proactive, e.g., LW), or high irreplaceability (e.g., G200). However, how these prioritization frameworks apply to global tree diversity and their respective effectiveness for tree conservation remains an open question despite the ecological and societal importance of trees.

We selected three frameworks of global biodiversity conservation prioritizations(*25*), namely the BH ((*51*) and Conservation International), the G200 (*52*), and LW (*111*). BH is defined as regions containing at least 1,500 vascular plants as endemics (standing for irreplaceable) whilst having less than 30% of its original natural habitats remaining (https://www.conservation.org/priorities/biodiversity-hotspots). Globally 36 biogeographically similar aggregations of ecoregions are proposed, which represent 2.4% of Earth’s land surface, and support 77% of the world’s plant endemics, and nearly 43% of endemic tetrapods (bird, mammal, reptile, and amphibian species) (*112*). G200 (World Wildlife Fund) is a suite of 238 ecoregions within biomes characterized by high species richness, endemism, taxonomic uniqueness, unusual ecological or evolutionary phenomena, or global rarity of major habitat type (irreplaceability or distinctiveness) (*52*), and includes 142 terrestrial, 53 freshwater, and 43 marine ecoregions. The LW is the 10% area with the lowest human pressure within each of the Earth’s 60 biogeographic realms with human pressure measured with an aggregate index, human footprint index, of human density, land transformation, access, and infrastructure (*111*). Each conservation priority framework represented one kind of the irreplaceability/vulnerability framework of systematic conservation planning, i.e., BH prioritizes low vulnerability (purely reactive: prioritizing areas of low threat but high irreplaceability), LW prioritizes high vulnerability (proactive: prioritizing areas of high threat and high irreplaceability), and G200 prioritizes high irreplaceability. The BH data layer was downloaded from (*113*); the upgraded LW data layer was obtained from (*114*), and the G200 terrestrial ecoregions layer was from the World Wildlife Fund (https://www.worldwildlife.org/publications/global-200). We aggregated all the three spatial layers to 110 × 110 km grid spatial resolution.

### Human pressure data

We used the recently proposed Human Modification map (*44*) as a proxy of human pressure. Compared to the commonly-used human footprint map (*111, 115*), human modification map was modelled with the incorporation of 13 most recent global-scale anthropogenic layers (with a median year of 2016) to account for the spatial extent, intensity, and co-occurrence of human activities, many of which showing high directly or indirectly impact on biodiversity (*30*). Human modification index values were extracted at a resolution of 1 km^2^. Based on the definition of (*44*), we categorized the human modification index into three groups representing different intensity levels of human pressure: low (values ≤0.1; e.g., Southwest Amazon moist forests); moderate (0.1 – 0.4; e.g., Everglades flooded grasslands); and high to very high (>0.4; e.g., Central Indochina dry forests). We plotted this human pressure layer with the top 17% and 50% priority areas to visualize them spatially.

### Prioritization Analyses

We used the Zonation systematic conservation prioritization software (v.4, (*41, 116*)) to identify global priority areas for tree diversity in each of the three dimensions. Zonation is based on the principle of complementarity, i.e., balancing a set of biodiversity features to jointly achieve the most complete representation in a given region, and evaluating the spatial priority areas through the priority ranking (*41*). We used the Core-Area Zonation (CAZ) algorithm as cell removal rule. The CAZ ranking algorithm emphasizes species (or any biodiversity feature) rarity to ensure high-quality locations for all features, even if these features occur in species-poor or expensive areas (*40, 42*).

We ran the Zonation spatial conservation prioritization procedure on species, phylogenetic groups, and traits separately to compare the mismatch or congruence between the resulting priority areas. We focused on each of the obtained priority rankings on both the top 17% (CBD 2020) and 50% (CBD 2050) of the highest conservation value areas (i.e., the cells with ranking values greater than or equal to 0.83 or 0.50).

However, combining the top priority areas of the three diversity dimensions lead to total greater areas to be protected than the 17% and 50% targets, namely 26.3% and 61.5%, respectively (Fig. 1). Thus, to obtain prioritizations made jointly across the three diversity dimensions, but consistent with the two area targets, we ran a joint analysis using the three diversity component (species, phylogenetic groups, traits) layers as input of the Zonation, and selected the top 17% and top 50% priority areas for further analyses. In addition, as a sensitivity analysis we also compared the resulting priority areas to those from simply combining the priority areas from the prioritizations on the separate diversity dimensions. We found the two analyses generally generated similar results, i.e., similar percentages of overlap with existing PAs, and conservation priority frameworks (Fig. 2 *vs*. Fig. S9), and similar human pressure of the priority areas inside and outside existing PAs (Fig. 4 *vs*. Fig. S10). Thus, we only report the joint results in the main text, i.e., given that only these adhere to the 17% and 50% protected area goals.

### Analysis of spatial data

We evaluated the degree of congruence of the top 17% and 50% priority areas generated from the three dimensions using the Venn diagram. Then we ran global gap analyses for the top priority areas, PAs and each of the global biodiversity conservation priority frameworks, by overlaying the three layers (i.e., the top priority areas, PAs, and each conservation priority framework) to calculate the level of protection in PAs, potential protection in the priority framework, and shared protection of the two global high priority areas (both 17% and 50%).

We divided the priority areas based on the ranking scores into three categories: areas with priority scores higher than 0.83 (top 17%), between 0.50 and 0.83 (top 17 -50%), and less than 0.50 (<50%). For areas within each category, we further separated them into two parts based on whether or not they are located inside or outside PAs. We then calculated the human modification index for each of the six regions to test how different the human pressure was. In addition, we computed the mean and median values of the human modification index for regions inside and outside PAs globally. One-way ANOVA was used to statistically test the differences between groups of human modification index values.

Despite the unprecedented coverage and quality of our tree species dataset, we are aware of certain limitations of the data such as missing data in many parts of India and functional traits. Although phylogenetic and functional diversity are usually highly related (*27, 86*), and the imputation of functional traits using the phylogenetic eigenvectors could increase the intensity of the relation, we found distinct priority areas from each dimension. Nonetheless, more data on the three aspects are needed to improve the data completeness.

## General

We thank Brad Boyle for valuable database and informatics assistance and advice, and TRY contributors for sharing their data. This work was conducted as a part of the BIEN Working Group, 2008–2012. We thank all the data contributors and numerous herbaria who have contributed their data to various data compiling organizations (see the Supplementary Materials) for the invaluable data and support provided to BIEN. We thank the New York Botanical Garden; Missouri Botanical Garden; Utrecht Herbarium; the UNC Herbarium; and GBIF, REMIB, and SpeciesLink. The staff at CyVerse provided critical computational assistance.

We acknowledge the herbaria that contributed data to this work: A, AAH, AAS, AAU, ABH, ACAD, ACOR, AD, AFS, AK, AKPM, ALCB, ALTA, ALU, AMD, AMES, AMNH, AMO, ANGU, ANSM, ANSP, AQP, ARAN, ARIZ, AS, ASDM, ASU, AUT, AV, AWH, B, BA, BAA, BAB, BABY, BACP, BAF, BAFC, BAI, BAJ, BAL, BARC, BAS, BBB, BBS, BC, BCMEX, BCN, BCRU, BEREA, BESA, BG, BH, BHCB, BIO, BISH, BLA, BM, BOCH, BOL, BOLV, BONN, BOON, BOTU, BOUM, BPI, BR, BREM, BRI, BRIT, BRLU, BRM, BSB, BUT, C, CALI, CAN, CANB, CANU, CAS, CATA, CATIE, CAY, CBM, CDA, CDBI, CEN, CEPEC, CESJ, CGE, CGMS, CHAM, CHAPA, CHAS, CHR, CHSC, CIB, CICY, CIIDIR, CIMI, CINC, CLEMS, CLF, CMM, CMMEX, CNPO, CNS, COA, COAH, COCA, CODAGEM, COFC, COL, COLO, CONC, CORD, CP, CPAP, CPUN, CR, CRAI, CRP, CS, CSU, CSUSB, CTES, CTESN, CU, CUVC, CUZ, CVRD, DAO, DAV, DBG, DBN, DES, DLF, DNA, DPU, DR, DS, DSM, DUKE, DUSS, E, EA, EAC, EAN, EBUM, ECON, EIF, EIU, EMMA, ENCB, ER, ERA, ESA, ETH, F, FAA, FAU, FAUC, FB, FCME, FCO, FCQ, FEN, FHO, FI, FLAS, FLOR, FM, FR, FRU, FSU, FTG, FUEL, FULD, FURB, G, GAT, GB, GDA, GENT, GES, GH, GI, GLM, GMDRC, GMNHJ, GOET, GRA, GUA, GZU, H, HA, HAC, HAL, HAM, HAMAB, HAO, HAS, HASU, HB, HBG, HBR, HCIB, HEID, HGM, HIB, HIP, HNT, HO, HPL, HRCB, HRP, HSC, HSS, HU, HUA, HUAA, HUAL, HUAZ, HUCP, HUEFS, HUEM, HUFU, HUJ, HUSA, HUT, HXBH, HYO, IAA, IAC, IAN, IB, IBGE, IBK, IBSC, IBUG, ICEL, ICESI, ICN, IEA, IEB, ILL, ILLS, IMSSM, INB, INEGI, INIF, INM, INPA, IPA, IPRN, IRVC, ISC, ISKW, ISL, ISTC, ISU, IZAC, IZTA, JACA, JBAG, JBGP, JCT, JE, JEPS, JOTR, JROH, JUA, JYV, K, KIEL, KMN, KMNH, KOELN, KOR, KPM, KSC, KSTC, KSU, KTU, KU, KUN, KYO, L, LA, LAGU, LBG, LD, LE, LEB, LIL, LINC, LINN, LISE, LISI, LISU, LL, LMS, LOJA, LOMA, LP, LPAG, LPB, LPD, LPS, LSU, LSUM, LTB, LTR, LW, LYJB, LZ, M, MA, MACF, MAF, MAK, MARS, MARY, MASS, MB, MBK, MBM, MBML, MCNS, MEL, MELU, MEN, MERL, MEXU, MFA, MFU, MG, MGC, MICH, MIL, MIN, MISSA, MJG, MMMN, MNHM, MNHN, MO, MOL, MOR, MPN, MPU, MPUC, MSB, MSC, MSUN, MT, MTMG, MU, MUB, MUR, MVFA, MVFQ, MVJB, MVM, MW, MY, N, NA, NAC, NAS, NCU, NE, NH, NHM, NHMC, NHT, NLH, NM, NMB, NMNL, NMR, NMSU, NSPM, NSW, NT, NU, NUM, NY, NZFRI, O, OBI, ODU, OS, OSA, OSC, OSH, OULU, OWU, OXF, P, PACA, PAMP, PAR, PASA, PDD, PE, PEL, PERTH, PEUFR, PFC, PGM, PH, PKDC, PLAT, PMA, POM, PORT, PR, PRC, PRE, PSU, PY, QCA, QCNE, QFA, QM, QRS, QUE, R, RAS, RB, RBR, REG, RELC, RFA, RIOC, RM, RNG, RSA, RYU, S, SACT, SALA, SAM, SAN, SANT, SAPS, SASK, SAV, SBBG, SBT, SCFS, SD, SDSU, SEL, SEV, SF, SFV, SGO, SI, SIU, SJRP, SJSU, SLPM, SMDB, SMF, SNM, SOM, SP, SPF, SPSF, SQF, SRFA, STL, STU, SUU, SVG, TAES, TAI, TAIF, TALL, TAM, TAMU, TAN, TASH, TEF, TENN, TEPB, TEX, TFC, TI, TKPM, TNS, TO, TOYA, TRA, TRH, TROM, TRT, TRTE, TU, TUB, U, UADY, UAM, UAMIZ, UB, UBC, UC, UCMM, UCR, UCS, UCSB, UCSC, UEC, UESC, UFG, UFMA, UFMT, UFP, UFRJ, UFRN, UFS, UGDA, UH, UI, UJAT, ULM, ULS, UME, UMO, UNA, UNB, UNCC, UNEX, UNITEC, UNL, UNM, UNR, UNSL, UPCB, UPEI, UPNA, UPS, US, USAS, USF, USJ, USM, USNC, USP, USZ, UT, UTC, UTEP, UU, UVIC, UWO, V, VAL, VALD, VDB, VEN, VIT, VMSL, VT, W, WAG, WAT, WELT, WFU, WII, WIN, WIS, WMNH, WOLL, WS, WTU, WU, XAL, YAMA, Z, ZMT, ZSS, and ZT.

## Funding

WYG, JMSD and JCS acknowledge support from the Danish Council for Independent Research | Natural Sciences (Grant 6108-00078B) to the TREECHANGE project. JCS also considers this work a contribution to his VILLUM Investigator project “Biodiversity Dynamics in a Changing World” funded by VILLUM FONDEN. CB was supported by the National Research Foundation of Korea (NRF) grant funded by the Korean government (MSIT) (2018R1C1B6005351). ASM was supported by the Environment Research and Technology Development Fund (S-14) of the Ministry of the Environment, Japan. JP (Jan Pisek) was supported by the Estonian Research Council grant PUT1355. JP (Josep Peñuelas) was funded by the European Research Council Synergy grant ERC-2013-SyG-610028 IMBALANCE-P. AGG (Alvaro G. Gutiérrez) was funded by FONDECYT 11150835 and 1200468.

The BIEN working group was supported by the National Center for Ecological Analysis and Synthesis, a center funded by NSF EF-0553768 at the University of California, Santa Barbara and the State of California. Additional support for the BIEN working group was provided by iPlant/CyVerse via NSF DBI-0735191. B.J.E. and C.M. were supported by NSF ABI-1565118 and NSF HDR-1934790. B.J.E. was also supported by the Global Environment Facility SPARC project grant (GEF-5810). B.J.E., C.V., and B.S.M. acknowledge the FREE group funded by the synthesis center CESAB of the French Foundation for Research on Biodiversity (FRB) and EDF. J.-C.S. and B.J.E. acknowledge support from the Center for Informatics Research on Complexity in Ecology (CIRCE), funded by the Aarhus University Research Foundation under the AU Ideas program.

## Author contributions

WYG, JMSD and JCS conceived the project; JMSD, WYG, and all others collected the data; WYG analyzed data with the help of KG and WBX; WYG interpreted the data; WYG, JMSD and JCS wrote the manuscript. All authors contributed data, discussed the results, revised manuscript drafts, and contributed to writing the final manuscript.

## Competing interests

The authors declare no competing interests.

## Data and materials availability

All the occurrences are deposited in BIEN (https://bien.nceas.ucsb.edu/bien/), and the phylogeny and imputed trait data are available as Supplementary data.

## Notes

### Competing Interest Statement

The authors have declared no competing interest.

