## Supplementary material for "Half of the world’s tree biodiversity is unprotected and is increasingly threatened by human activities": Supplemantary material

### Supplementary Materials

**Table S1** Pearson correlation coefficients (all significant at *p* level of 0.05) between the species richness, phylogenetic diversity, functional diversity, conservation priority for taxonomic dimension, conservation priority for phylogenetic dimension, and conservation priority for functional trait dimension.

| | taxonomic dimension | phylogenetic dimension | trait dimension |
|---|---|---|---|
| taxonomic dimension | 1 | 0.766 | 0.763 |
| phylogenetic dimension | 0.766 | 1 | 0.982 |
| trait dimension | 0.763 | 0.982 | 1 |

**Table S2** The 21 functional traits compiled and their completeness in the 54020 species included in the phylogeny. At the imputation, eight traits were selected to compute the functional trait diversity.

| Trait name | Description | Missing species (totally 54,020 species) | Missing rate (%) | Selected |
|---|---|---|---|---|
| Leaf nitrogen content | Leaf nitrogen (N) content per leaf dry mass | 50,441 | 93.37 | √ |
| Wood Density | Stem specific density (SSD) | 47,608 | 88.13 | √ |
| Leaf K conent | Leaf potassium (K) content per leaf dry mass | 52,778 | 97.70 | |
| Leaf P content | Leaf phosphorus (P) content per leaf dry mass | 51,652 | 95.62 | √ |
| LDMC | Leaf dry matter content | 53,227 | 98.53 | √ |
| Vegetative height | Plant maximum height | 49,521 | 91.67 | √ |
| Leaf N:P | Leaf nitrogen/phosphorus (N/P) ratio | 52,566 | 97.31 | |
| Seed dry mass | Seed dry mass | 49,348 | 91.35 | √ |
| SLA | specific leaf area (Leaf area per leaf dry mass, 1/LMA) | 52,055 | 96.36 | √ |
| LA | Leaf area (in case of compound leaves: leaflet, petiole and rachis excluded) | 53,221 | 98.52 | √ |
| SLAFM | Leaf area per leaf fresh mass (SLA based on leaf fresh mass) | 53,739 | 99.48 | |
| $LA_{compundleaf}$ | Leaf area (in case of compound leaves: leaf, petiole included) | 53,355 | 98.77 | |
| Leaf respiration rate | Leaf respiration rate per leaf area | 53,802 | 99.60 | |
| Wood N | Wood nitrogen (N) content per wood dry mass | 53,989 | 99.94 | |
| Leaf photosynthesis rate | Leaf photosynthesis rate per leaf area | 53,252 | 98.58 | |
| Seed germination rate | Seed germination rate (germination efficiency) | 53,339 | 98.74 | |
| LWR | Leaf dry mass per plant dry mass (leaf weight ratio) | 53,828 | 99.64 | |

| | | | |
|---|---|---|---|
| Stomata conductance | Stomata conductance per leaf area | 53,542 | 99.12 |
| Fine root DM | Fine root dry mass to leaf dry mass ratio | 53,948 | 99.87 |
| Leaf WUE | Leaf photosynthetic water use efficiency | 53,970 | 99.91 |
| Wood DM | Wood dry mass per plant | 54,012 | 99.99 |

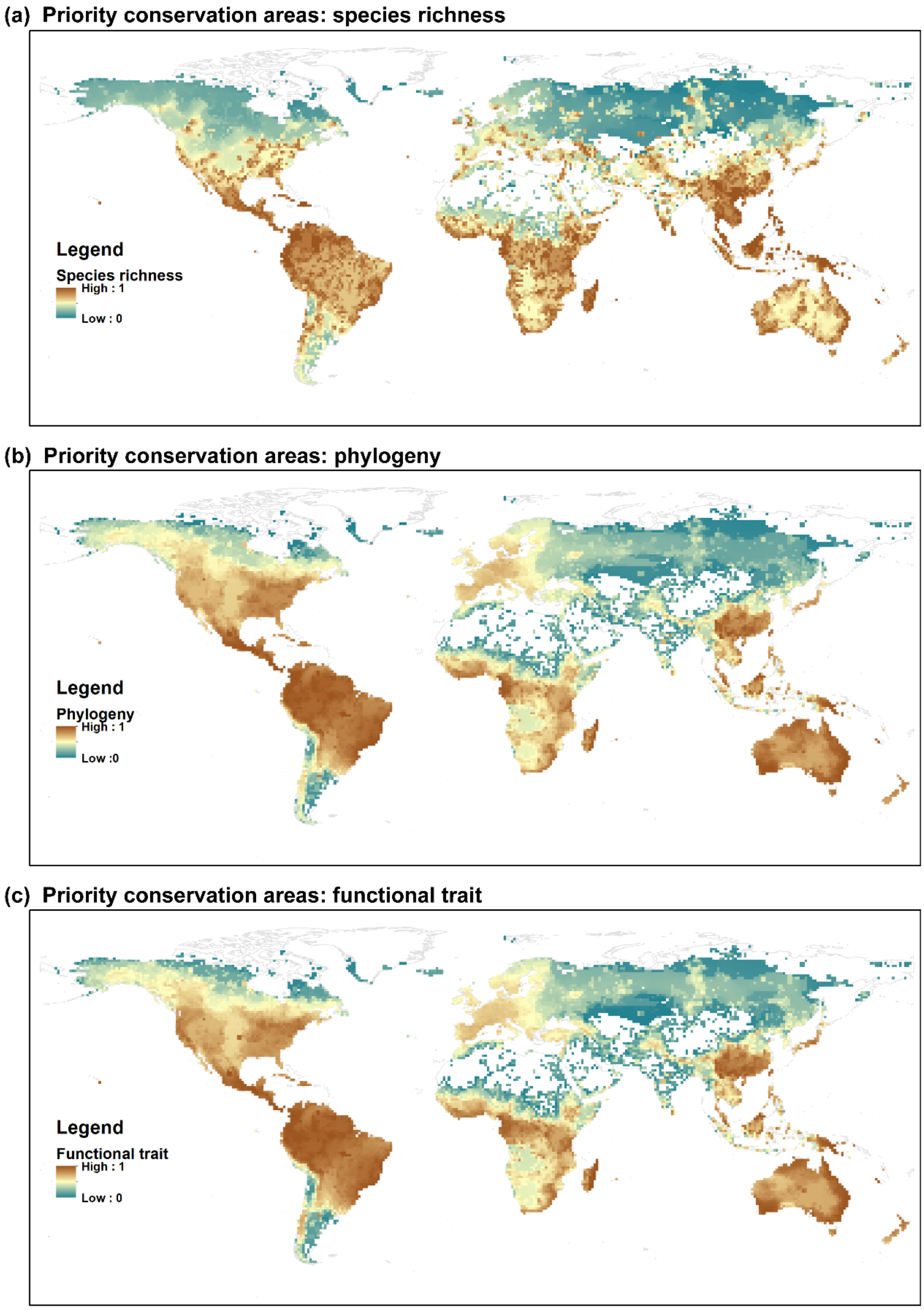

**Fig. S1** The priority conservation areas showing the ranking from each of the three dimension analyses. (a) Species richness, (b) Phylogeny, and (c) Functional traits. The values of cell rank from 0 to 1 with the increase of the conservation importance.

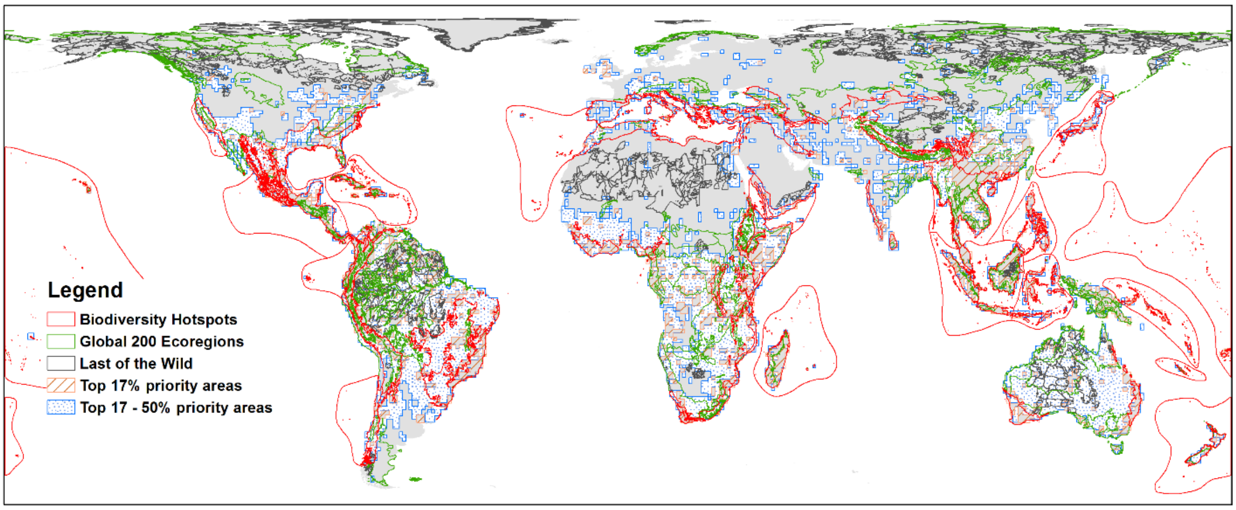

**Fig. S2** Maps showing the top 17% (CBD 2020 target) and top 50% (CBD 2050 target) priority areas obtained in the study, Global 200 Ecoregions (G200), the Last of the Wild (LW), and the Biodiversity Hotspots (BH).

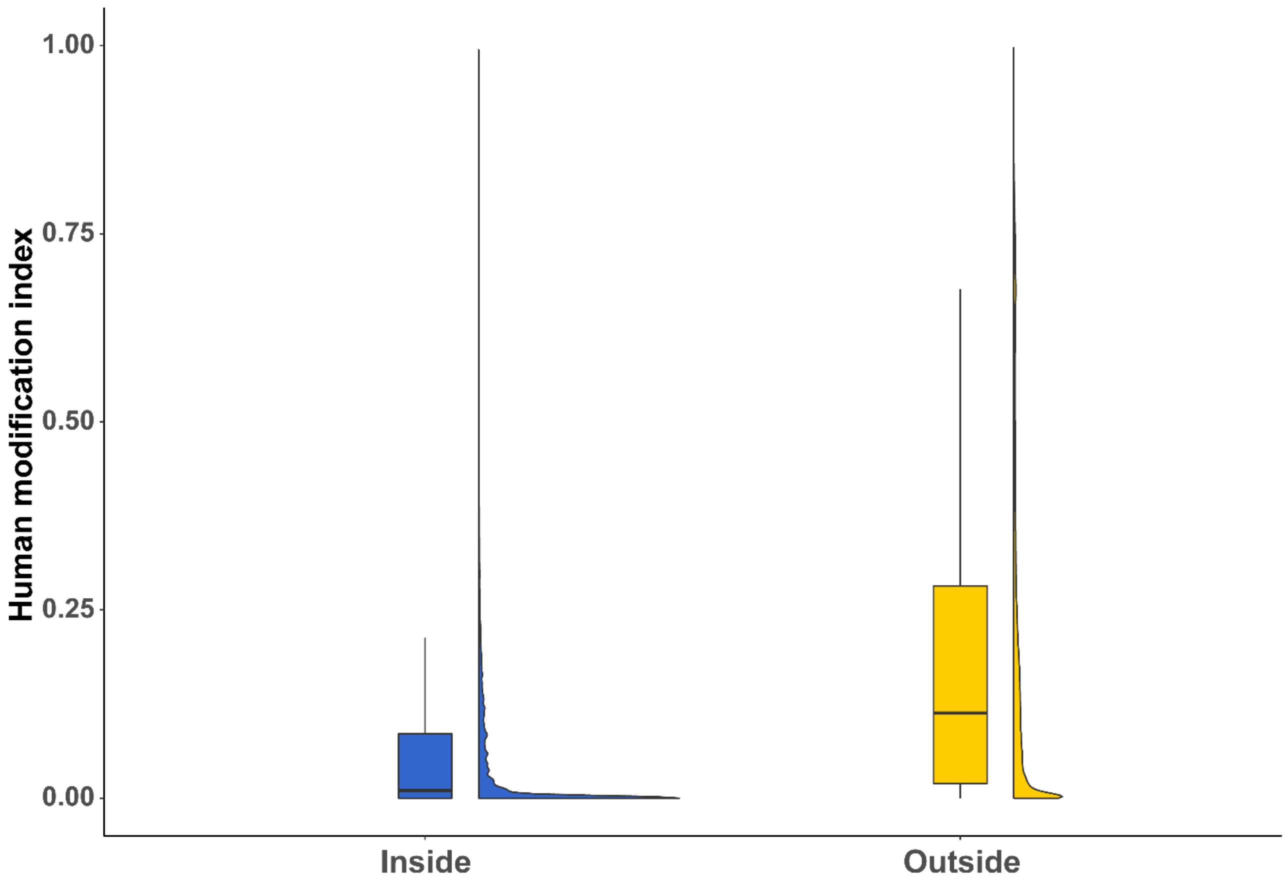

**Fig. S3** Boxplot and density plot of the global human modification index inside and outside existing PAs. One-way ANOVA test: F-value = 4531569, $p < 0.0001$. Means are indicated by circles in the boxplots. Sample size of the inside and outside PAs is 8,577,868 and 125,609,765, respectively.

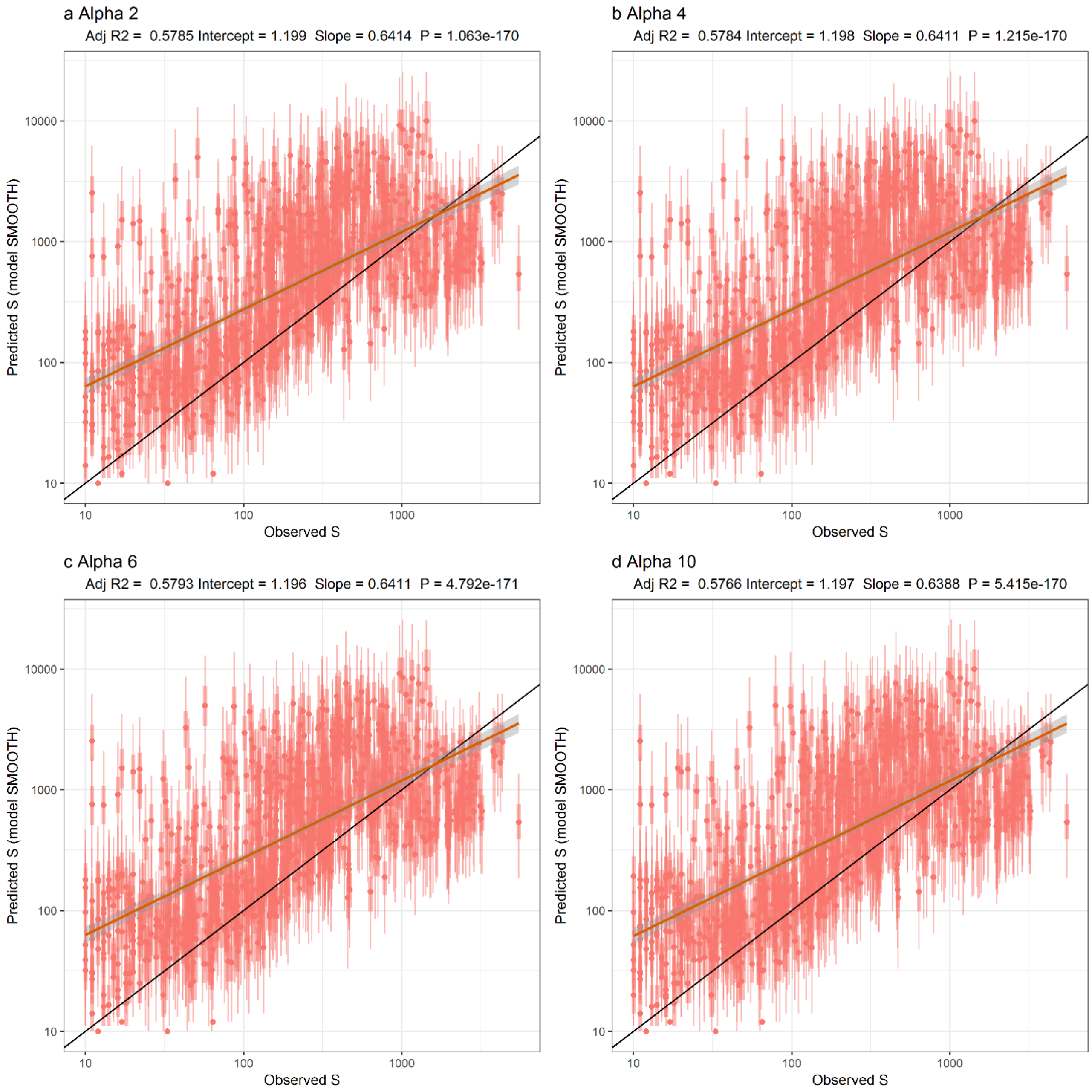

**Fig. S4** External validation of the observed richness (S) from each alpha-value range map using the predictions of model SMOOTH at the grain of the 209,903 $km^2$ hexagons (Keil & Chase 2019). Each subplot shows how the observed S compares with predictions of model SMOOTH from Keil & Chase (2019). Provided are full Bayesian prediction intervals, reflecting both the uncertainty in model parameters, as well as the width of the negative binomial error distribution. Points are medians of posterior distributions of the predictions; thick transparent bars are 25% and 75% quantiles, thin lines are 2.5% and 97.5% quantiles of the posterior distributions. Solid diagonal line is 1:1 line, and solid red line is the regression line between the observed and predicted S, with the confidence intervals in shade. The model performance is provided above each subplot.

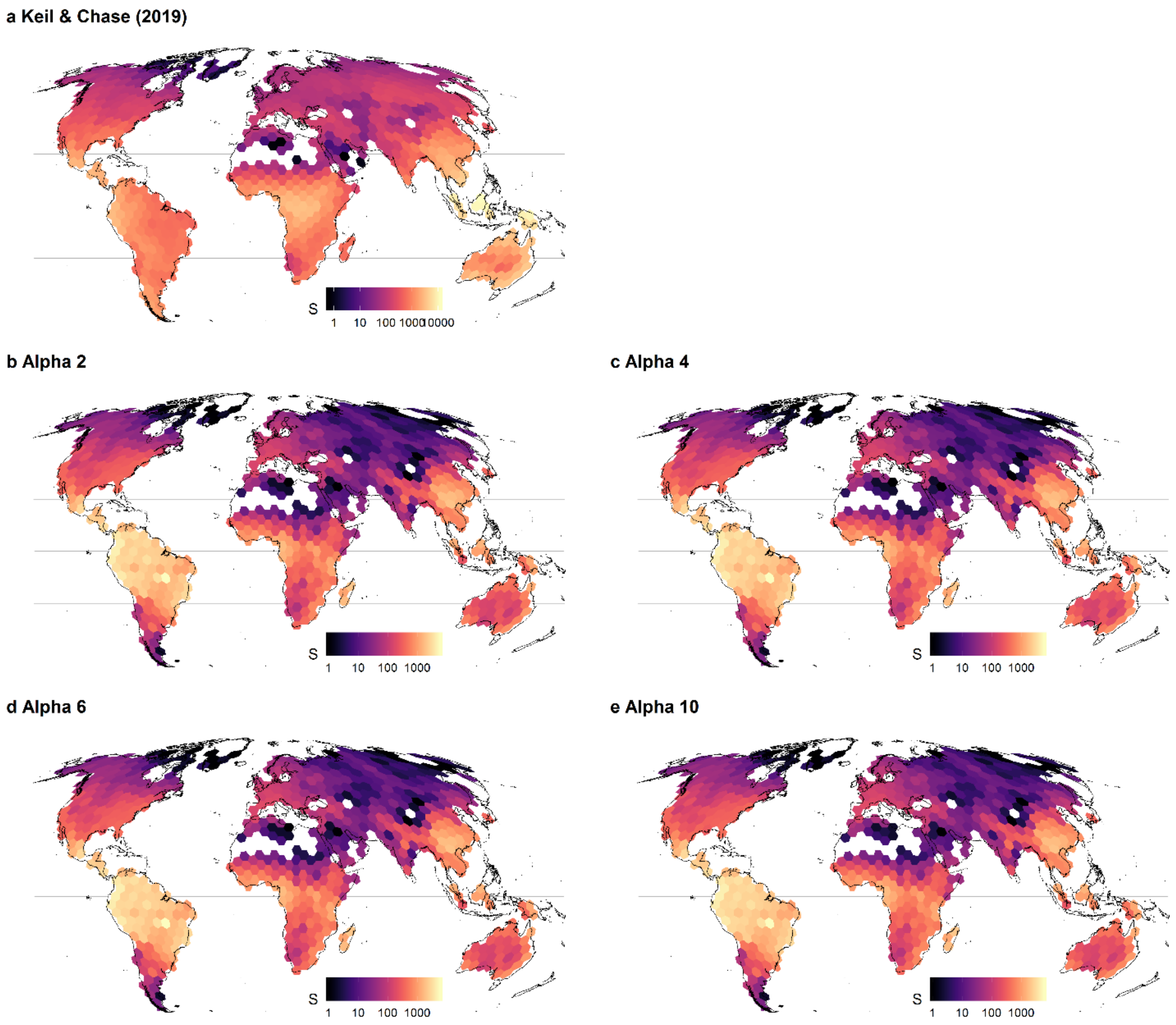

**Fig. S5** Tree-species richness (S) at the grain of the 209,903 km$^2$ hexagons. (a) Predicted pattern from model SMOOTH in Keil & Chase (2019); (b-e), observed patterns of the alpha-level range maps at the same grain size. White hexagons represent data missing or S = 0.

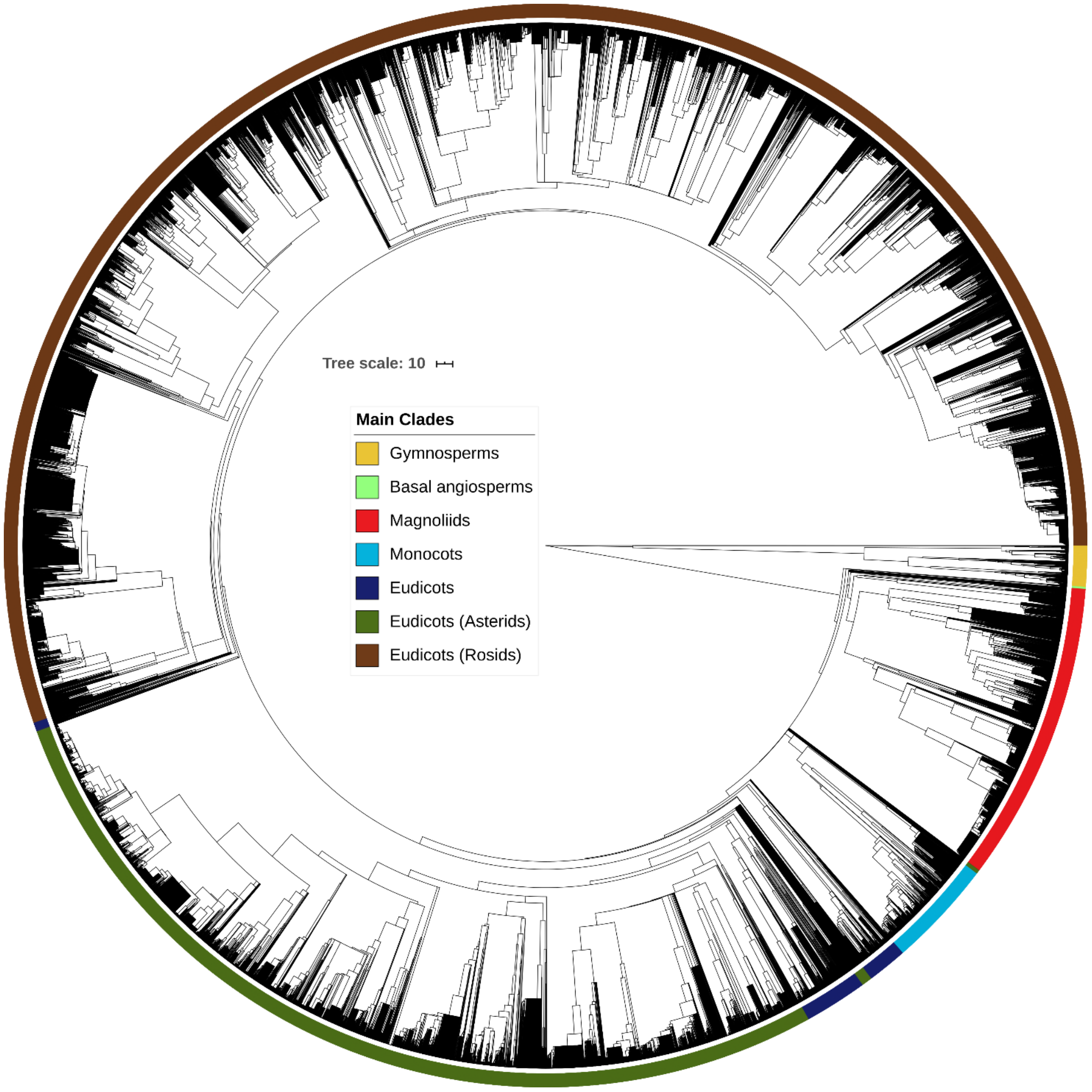

**Fig. S6** Time-calibrated phylogeny of the 46,752 tree species included in the study. Colors indicate main clades.

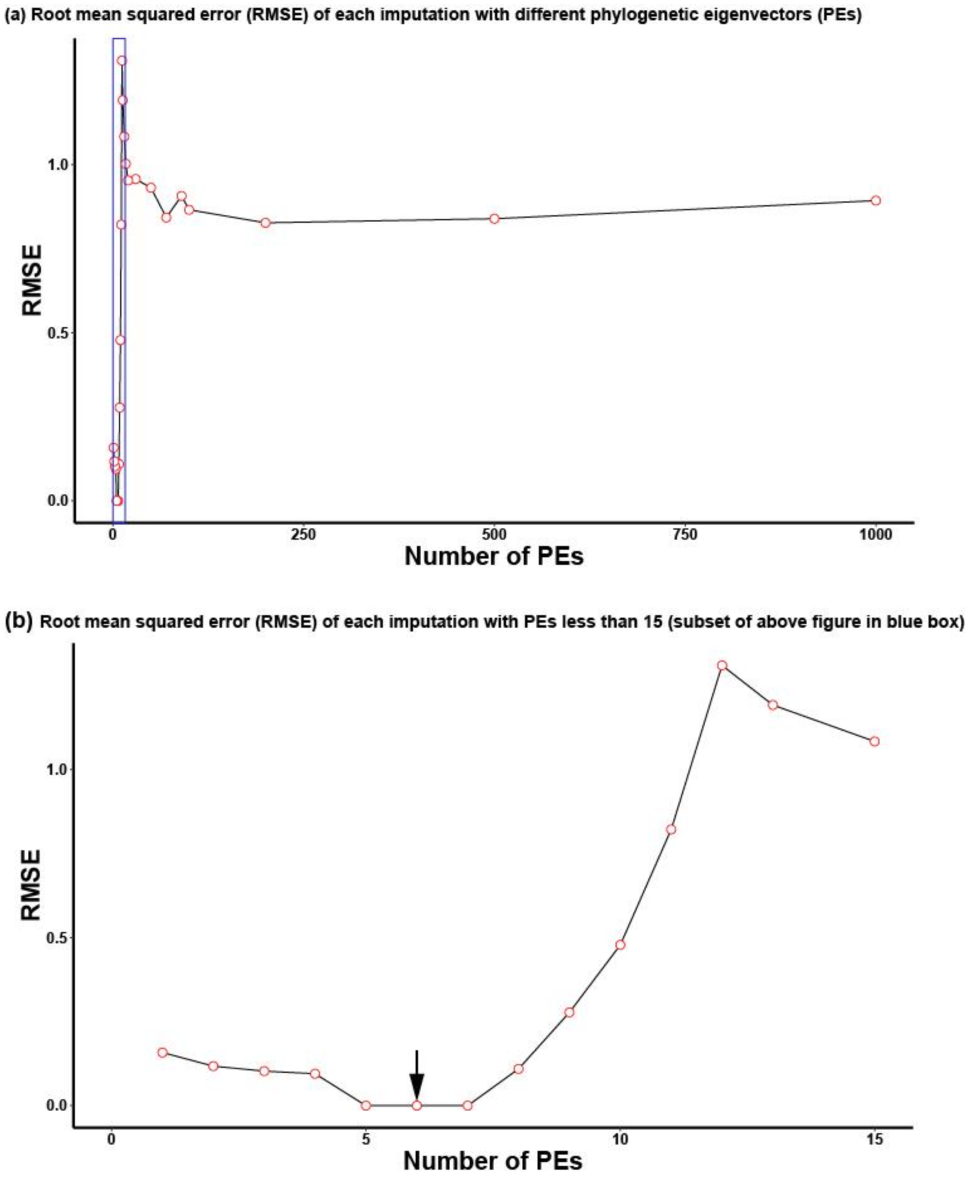

**Fig. S7** Root mean squared errors (RMSEs) from each Bayesian Hierarchical Probabilistic Matrix Factorization (BHPMF) imputation using different numbers of phylogenetic eigenvectors (PEs). RMSE is a quadratic scoring rule that also measures the average magnitude of the error; it is the square root of the average of squared differences between prediction and actual observation. It can range from zero to $\infty$ and is indifferent to the direction of errors. It is negatively oriented scores, which means lower values are better. The RMSE is the biggest for the imputation with 12 PEs, then it decreases with increasing PEs, and keeps stable at about 0.85 until the imputation with 1000 PEs (the maximum number of PEs investigated) (a). (b) The imputation with six PEs has the smallest RMSE value (0.087), followed by imputations with five or seven PEs (0.089). Thus, the final imputation selected six PEs (the black arrow in (b), which is an enlarged figure to show the part in the blue box in (a)).

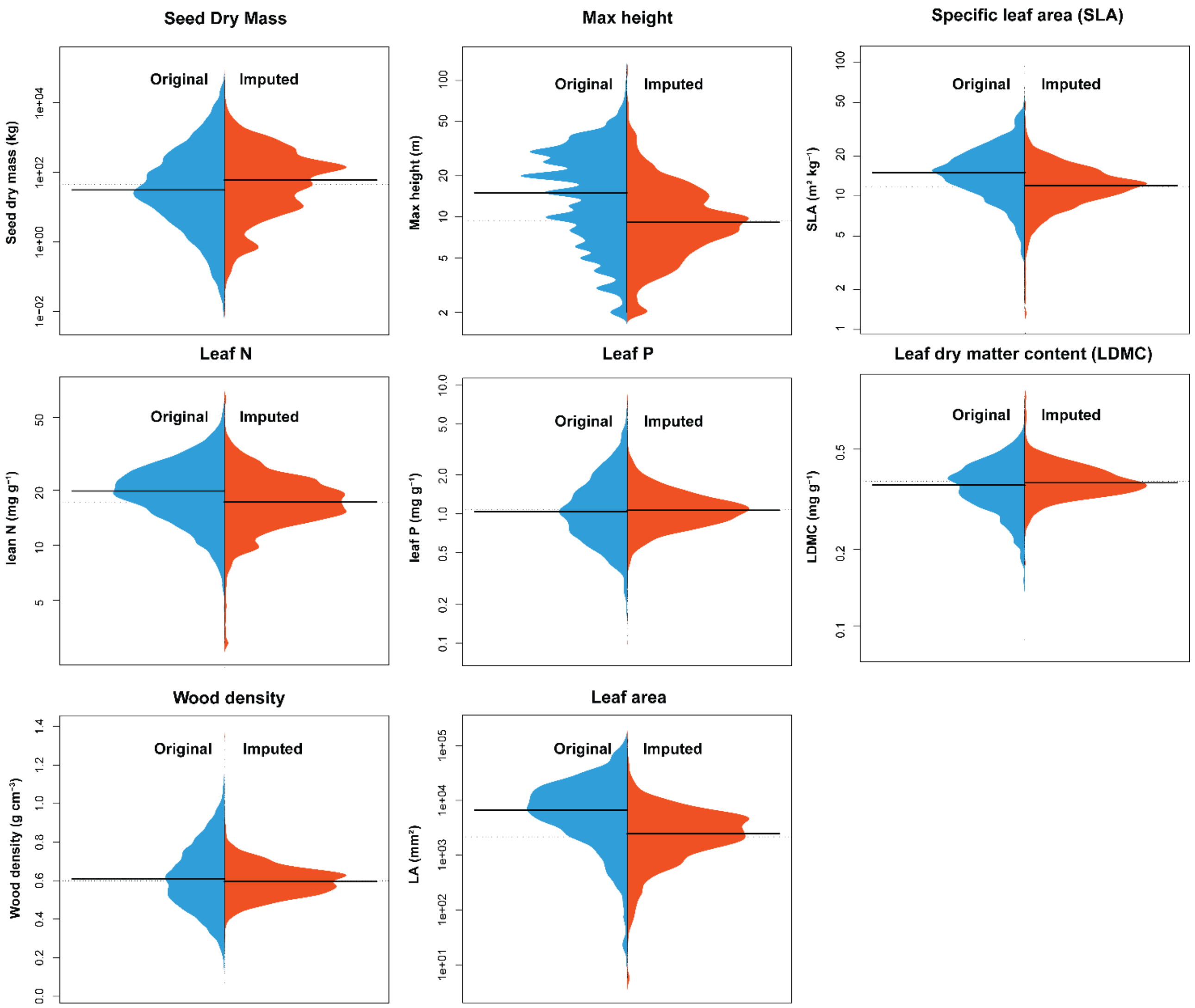

**Fig. S8** Asymmetric beanplots to compare the density distributions of original and imputed data for the eight functional traits used to calculate functional diversity. The black line indicates the median of each dataset.

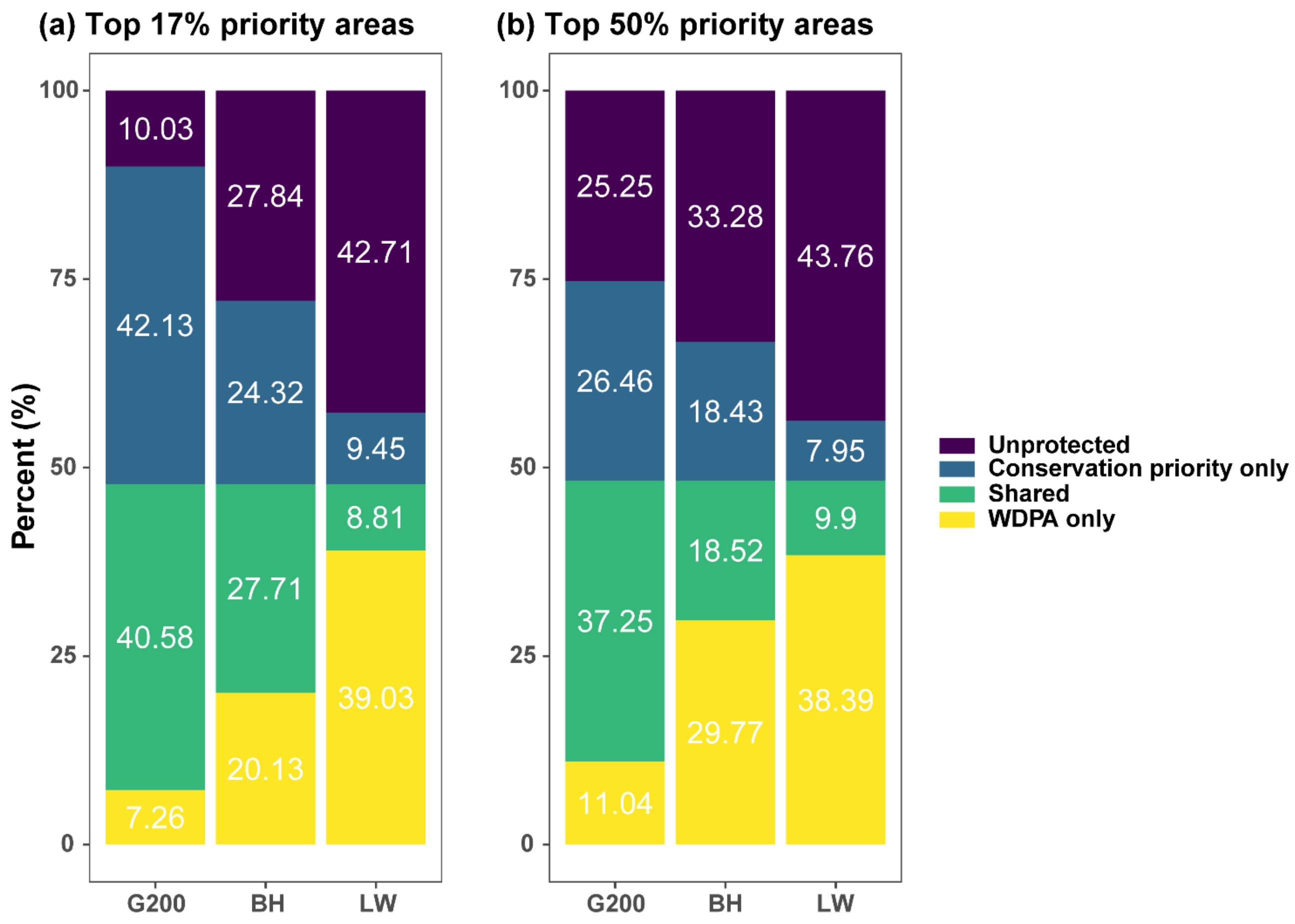

**Fig. S9** Percentages of the top 17% (CBD 2020 target) and top 50% (CBD 2050 target) priority areas for tree diversity covered by the existing protected areas (WDPA) or by each global biodiversity conservation priority framework (G200, BH, and LW). Data were from the combined priority areas, i.e., combining the top priority areas of the three diversity dimensions. Unprotected: areas not overlapping with either WDPA or the conservation priority framework; Conservation priority only: areas overlapping with conservation priority framework only; Shared: areas overlapping with both WDPA and conservation priority framework; WDPA only: areas overlapping with WPDA only. G200: Global 200 Ecoregions; BH: Biodiversity Hotspots; LW: Last of the Wild.

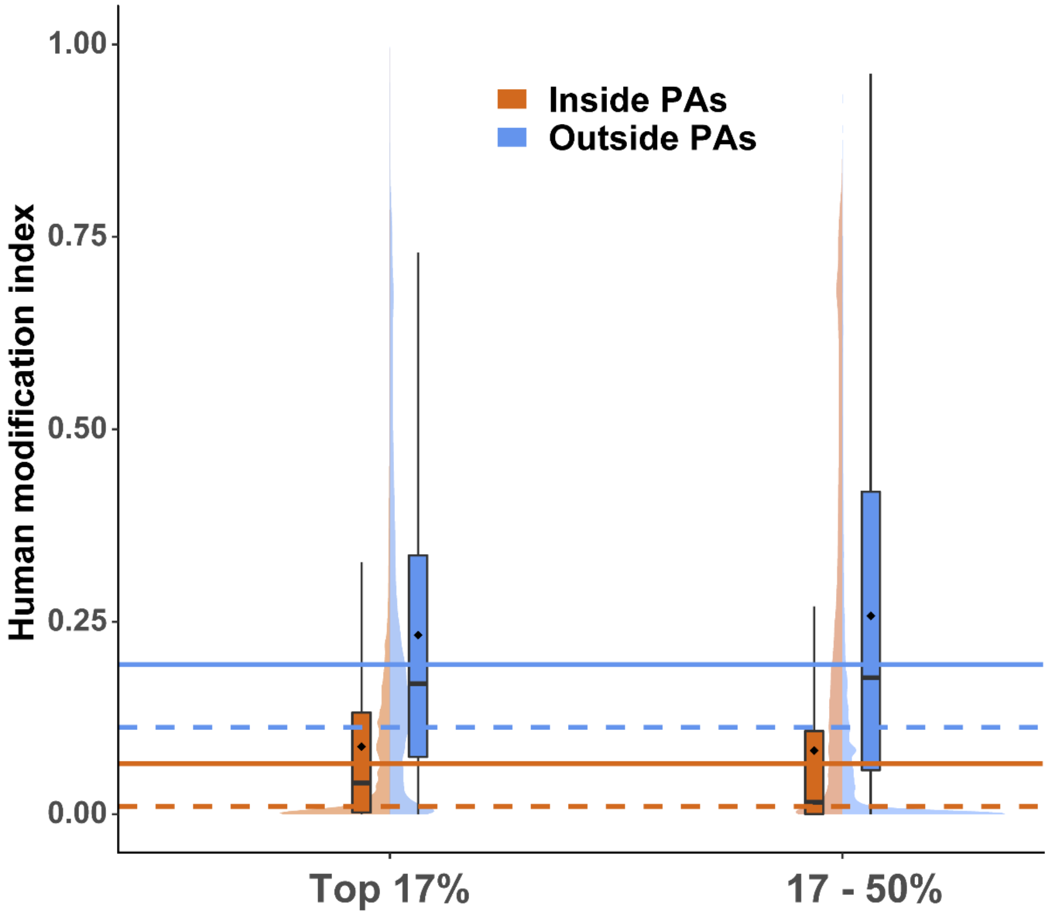

**Fig. S10** Split violin density plot of the human modification index across tree diversity conservation categories (top 17%, and top 17 - 50% of the priority scores from the Zonation prioritization). Data were from the combined priority areas, i.e., combining the top priority areas of the three diversity dimensions. Each category is divided into two groups: areas inside (dark orange) and outside (light blue) protected areas (PAs, following the World Database on Protected Areas (WDPA) database). For each category, there are two subplots: a boxplot and density plot. Medians and means each are indicated by solid lines and points in the boxplots. The horizontal solid lines and dashed lines indicate the global mean and median human modification values for the areas inside and outside PAs, respectively. All values were obtained from 1- $km^2$ resolution input layers (see Methods for more information).
